# Dysregulated calcium signaling in the aged macaque entorhinal cortex associated with tau hyperphosphorylation

**DOI:** 10.1101/2024.12.05.626721

**Authors:** S Bathla, D Datta, D Bolat, E Woo, A Duque, J Arellano, AFT Arnsten, AC Nairn

## Abstract

Tau pathology in sporadic Alzheimer’s disease (AD) follows a distinct pattern, beginning in the entorhinal cortex (ERC) and spreading to interconnected brain regions. Early-stage tau pathology, characterized by soluble phosphorylated tau, is difficult to study in human brains post-mortem due to rapid dephosphorylation. Rhesus macaques, which naturally develop age-related tau pathology resembling human AD, provide an ideal model for investigating early tau etiology. This study examines the molecular processes underlying tau pathology in the macaque ERC, focusing on calcium and inflammatory signaling pathways. Our findings reveal age-related decreases in PDE4 phosphodiesterases that hydrolyze cAMP and increases in calpain-2 and GCPII that occur in parallel with early-stage tau hyperphosphorylation at multiple epitopes (pS214-tau, pT181-tau, pT217-tau). These findings suggest that dysregulated calcium signaling in ERC, beginning in middle-age, primes tau for hyperphosphorylation, potentially driving the early stages of AD, advancing our understanding of how ERC vulnerabilities contribute to neurodegeneration in AD.

## Introduction

Tau pathology in sporadic Alzheimer’s disease (AD) exhibits a stereotypical progression pattern with neurofibrillary tangles (NFTs) in cortex forming earliest in the rhinal cortices, especially in the entorhinal (ERC) and transentorhinal cortex (Braak and Braak 1991, Braak and Braak 1992, Braak and Del Trecidi 2015, Braak and Del Tredici 2015, Hyman et al. 1986, Kaufman et al. 2018). Notably, NFTs appear in middle age, distinctly in the layer II cell islands in the ERC (Braak and Del Tredici 2015, Liu and Li 2019). Evidence from studies in animals and humans indicates that tau pathology spreads from the ERC to interconnected glutamatergic neurons in the limbic and association cortices, as well as hippocampus, seeding pathology throughout higher brain circuits, with primary sensory cortices impacted only at end stage disease (Ahmed et al. 2014, Calafate et al. 2015, Colin et al. 2020, de Calignon et al. 2012, Dujardin et al. 2018, Fu et al. 2018, Kaufman, et al. 2018, Lewis et al. 1987). Therefore, illuminating why the ERC is highly susceptible to tau pathology is critical to uncovering the etiology of the common, late-onset, sporadic form of AD. However, the earliest stage, soluble phosphorylated tau cannot be studied in human brains except by using biopsy samples, as it is rapidly dephosphorylated post-mortem, within 10-15 minutes after death (Matsuo et al. 1994, Wang et al. 2015). Thus, preclinical animal models are needed to study the early etiology of tau pathology.

Rhesus macaques naturally develop tau pathology with advancing age, with the same *qualitative* features as human patients with AD, and thus can be utilized to understand why ERC circuits are especially vulnerable (Paspalas et al. 2018). Importantly, AT8-labeled (pS202/pT205-tau) neurofibrillary tangles can be seen in macaques of extreme age, with paired helical filaments identical to human (Paspalas, et al. 2018). The pattern and sequence of cortical tau pathology in macaques is also the same as in humans, first arising in the ERC layer II cell islands, and then extending into deeper ERC layers, the hippocampus, limbic and association cortices, with little expression in primary visual cortex (Arnsten, Datta, Del Tredici, et al. 2021, Arnsten et al. 2019, Arnsten, Datta and Preuss 2021, Paspalas, et al. 2018). Importantly, extremely short or virtually zero post-mortem intervals are possible when analyzing rhesus macaque brains, which allows the analysis of early, soluble forms of hyperphosphorylated tau. This includes the capture of tau phosphorylated at threonine 181 (pT181-tau) and threonine 217 (pT217-tau), tau species used as fluid-based biomarkers in humans, where pT217-tau in particular is being developed as a plasma biomarker for incipient AD (Barthelemy et al. 2020, Hansson et al. 2023, Janelidze et al. 2020, Mattsson-Carlgren et al. 2023, Olsson et al. 2016, Salvado et al. 2023).

Longstanding research has suggested that calcium dysregulation might be a crucial precipitating factor in AD pathogenesis (Alzheimer’s Association Calcium Hypothesis 2017, Gibson and Peterson 1987, Khachaturian 1994). Research from aging macaques has corroborated this hypothesis, showing that excitatory cells in vulnerable association cortices such as the ERC and the dorsolateral prefrontal cortex (dlPFC) express the molecular machinery for cAMP-PKA actions to magnify calcium signaling, particularly within dendrites and dendritic spines, and that dysregulation with age and/or inflammation contributes to tau hyperphosphorylation (Arnsten et al. 2020, Arnsten, Datta and Wang 2021, Arnsten and Wang 2020). For example, cAMP-calcium regulation by PDE4D and mGluR3 are reduced in the aged macaque dlPFC (Arnsten, Datta and Preuss 2021, Datta et al. 2020b). Elevated PKA signaling phosphorylates ryanodine receptors (pRyR2) on the SER to cause calcium leak into the cytosol, which is seen in the aged dlPFC and in middle age in the more vulnerable ERC (Datta et al. 2021, Paspalas, et al. 2018), and has been documented in the brains of patients with Alzheimer’s disease (Lacampagne et al. 2017). Very high levels of cytosolic calcium can activate calpain-2, which cleaves and activates GSK3β and p25-cdk5 contributing to the hyperphosphorylation of tau (Arnsten, Datta and Preuss 2021, Arnsten, Datta and Wang 2021, Baudry et al. 2013). However, it is not known if signs of dysregulated calcium signaling can be seen in association with hyperphosphorylated in the macaque ERC, where cortical tau pathology first begins. These relationships were explored in the current study of rhesus macaque ERC, utilizing both biochemistry and immunohistochemistry to examine molecular features of pathology with advancing age. We examined the expression patterns of early stage, soluble phosphorylated tau (pS214-tau, pT181-tau, pT217-tau), as well as the expression of mechanisms that regulate cAMP-calcium signaling (PDE4A, PDE4D, mGluR3), and those that drive pathology (S2808RyR2, calpain-2, GCPII) in rhesus macaque ERC across the adult age-span.

## Materials and Methods

Animals were cared for in accordance with the guidelines of Yale University Institutional Animal Care and Use Committee, and Public Health Service requirements for animal use as described in the Guide for the Care and Use of Laboratory Animals. Yale University is accredited by the American Association for Accreditation of Laboratory Animal Care (AAALAC).

### Animal and Tissue processing for biochemistry

Rhesus monkeys used for biochemical experiments ranged in age from 8.3 to 28.6 years (N = 10, all female). PMI was minimized to 10 min to ∼1 hr as longer PMI has significant impact on tissue quality and more specifically on levels of phosphorylation. For tissue collection, the dura was removed, and ERC tissue taken out using a scalpel. Immediately following dissection samples were placed into liquid nitrogen and stored at −80°C for further use.

### Protein Extraction

Brain tissue (ERC, 100 mg) was lysed in 1% Triton X-100 lysis buffer (200 mM NaCl, 10 mM HEPES, 10 mM EGTA, 10 mM EDTA, phosSTOP phosphatase inhibitor, and cOmplete mini protease inhibitor) with 20 strokes in a homogenizer. Cell debris was removed by centrifugation for 15 min (13,000 x g) at 4°C. The supernatant was collected, and protein concentration was determined with Bradford Assay (Bio-Rad, USA). The supernatant was stored at −80^0^C for further use.

### Immuno-blotting

Protein (40 ug per lane) was boiled for 5 min at 100°C in SDS-loading buffer with DTT. The samples were separated on 4-20% Tris-glycine gels using 150 V over 1.5 h in a Criterion cell (Bio-Rad, USA). Proteins were transferred onto 0.45 um nitrocellulose membranes at 300 mA for 1.5 hr in a Criterion blotter (Bio-Rad, USA). After 1 hr blocking at room temperature in TBST (20 mM Tris-HCl, 140 mM NaCl, pH 7.5, 0.05% Tween-20) containing 3% BSA, membranes were probed overnight with antibodies:pS214-tau (Abcam ab4846, 1:1000), pT181-tau (CST12885S,1:1000), pT217-tau antibody (AS-54968; AnaSpec, 1:1000), calpain 2 (Abcam ab39165,1:1000), PDE4A (Abcam ab14607,1:1000), PDE4D (Millipore ABS22, 1:1000), mGLUR3 (Abcam AB166608,1:1000), phospho-RyR S2808 (Abcam ab59225 1:500), GAPDH (Millipore CB1001-500, 1:10,000), GCPII (Proteintech, 13163-1-AP,1:1000) in TBST containing 3% BSA at 4°C. Membranes were washed 3 times with TBST and incubated with fluorescent secondary antibodies (1:10000) of the appropriate species for 1 hr at room temperature. Three TBST washes were used to remove secondary antibody. Blots were rinsed with Milli-Q water and analyzed using a LI-COR Odyssey scanner.

## Statistical analysis of biochemical data

Image Studio Lite was used for band quantification and background subtraction. Prior to quantification, background subtraction was done by calculating the average intensity immediately above and below the band(s) of interest. The expression of targets was further normalized by calculating the ratio of band intensity of target/band intensity GAPDH. For statistical comparison, unpaired t-test with Welch’s correction was performed on the grouped analysis as markers were normally distributed. The normalized data was plotted using Graphpad Prism software. The correlation analysis between age and targets (phosphorylated tau and Ca^+^ regulatory proteins) was determined by Pearson correlation.

### Immunohistochemistry

#### Animals and Tissue Preparation

Two young (8 and 10 years) and four aged (24, 28, 30 and 31 years) rhesus macaques (*Macaca mulatta*) were used for this study. As described previously (Datta et al. 2020a, Datta, et al. 2021, Jin et al. 2018b, Paspalas et al. 2013), rhesus macaques were deeply anesthetized prior to transcardial perfusion of 100 mM phosphate-buffer saline (PBS), followed by 4% paraformaldehyde/0.05% glutaraldehyde in 100 mM PBS. Following perfusion, a craniotomy was performed, and the entire brain was removed and dissected, including a frontal block containing the primary region of interest. The brains were sectioned coronally at 30 μm on a vibratome (Leica) across the entire rostrocaudal extent of the entorhinal cortex (ERC). The free-floating sections were cryoprotected in a solution containing ethylene glycol (30%), glycerol (30%) in 200 mM PB and stored at −20°C. The number of subjects was necessarily small, given the scarcity of macaques since their extensive use to develop SARS-cov-2 vaccines.

#### Histology and Immunoreagents

We used previously well-characterized primary antibodies raised in rabbit and mice. We used an affinity isolated polyclonal PDE4D protein (SAB4502128; Millipore Sigma Aldrich, Burlington, MA; RRID:AB_10744568) raised against amino acids 156-205 of PDE4D that recognizes human and rodent PDE4D based on sequence homology. The antibody is highly specific and detects endogenous levels of total PDE4D protein at a band migrating at ∼91 kDa. The antibody is suited for a range of applications, including immunohistochemistry, immunoblotting and ELISA as per manufacturer’s recommendations. The specificity and selectivity of the PDE4D antibody has been previously characterized using immunohistochemistry in myocytes to identify a role of PDE4D-PRKAR1α in cardiac contractility (Bedada et al. 2016) and with immunohistochemistry and immunoEM in rhesus macaque dlPFC (Datta, et al. 2020b). We used a rabbit anti-pT217-tau at 1:200 (cat# AS-54968, Anaspec). The immunogen used KLH conjugated with a synthetic phosphopeptide corresponding to human tau at phosphorylated threonine 217. We used a mouse anti-phosphoSer214-tau IgM (clone CP3) at 1:200 (generously provided by Dr. Peter Davies, The Feinstein Institutes for Medical Research) and a mouse anti-phosphoThr181-tau IgG1k (clone AT270) at 1:200 (MN1050; Thermo Fisher Scientific), antibodies that have been extensively validated by our group using immunohistochemistry and immunoelectron microscopy in rhesus macaque association cortices (Datta, et al. 2021, Paspalas, et al. 2018). For calpain-2, we used a rabbit anti-calpain-2 IgG at 1:200 (ab39165; Abcam). The immunogen is a synthetic peptide based on the amino terminal end of domain-III in the large subunit of calpain-2 that does not cross-react with other calpain family members and has been extensively validated in several protocols including immunohistochemistry, immunocytochemistry and immunofluorescence approaches.

#### Single-label immunoperoxidase immunohistochemistry

For single-label immunoperoxidase immunohistochemistry, sections of ERC were transferred for 1 hr to Tris-buffered saline (TBS) containing 5% bovine serum albumin, plus 0.05% Triton X-100 to block non-specific reactivity, and incubated in primary antibodies in TBS for 72 hr at 4°C. The tissue sections were incubated in goat anti-rabbit and goat anti-mouse biotinylated antibodies (Vector Laboratories) at 1:300 in TBS for 2 hr, and developed using the Elite ABC kit (Vector Laboratories) and diaminobenzidine (DAB) as a chromogen. Omission of the primary antibody eliminated all labeling. Sections were mounted on microscope slides and ERC cortical layers were photographed under an Olympus BX51 microscope equipped with a Zeiss AxioCam CCD camera. Zeiss AxioVision imaging software was used for imaging and data acquisition.

## Results

### Age-related increases in tau phosphorylation in rhesus macaque ERC

Biochemical analyses can accurately measure levels of soluble, phosphorylated tau. The current study assayed tau phosphorylated at S214, T217 and T181, three key early sites propelling tau pathology. We first quantified pS214-tau and observed significantly higher pS214-tau expression levels in aged monkeys compared to young (**Figure 1A-B**, \**P* =0.0012, t-5.277, df-6.074) in ERC. We found a trend-level positive correlation (\**P* = 0.0586) between the expression of pS214-tau and age in rhesus macaque ERC (**Figure 1C**). We further analyzed two additional phosphorylation sites (pT181-tau and pT217-tau) that have emerged as highly sensitive fluid-based biomarkers for the early identification of patients at risk of developing AD. pT181-tau expression was much higher in aged macaques than in younger animals (**Figure 2A-B**, *P =0.0004, t-6.412, df-6.988) in ERC. Furthermore, the protein expression of pT181-tau in rhesus macaque ERC were found to be positively correlated (**Figure 2C**, *P = 0.1779) with advanced age. Aged macaques also had significantly higher expression of pT217-tau (**Figure 3A-B**, *P =0.0235, t-3.003, df-6.074) compared to young animals in ERC, with a trend-level positive correlation between pT217-tau and age-span in rhesus macaque ERC (**Figure 3C**, *P = 0.2370).

**Figure 1:**
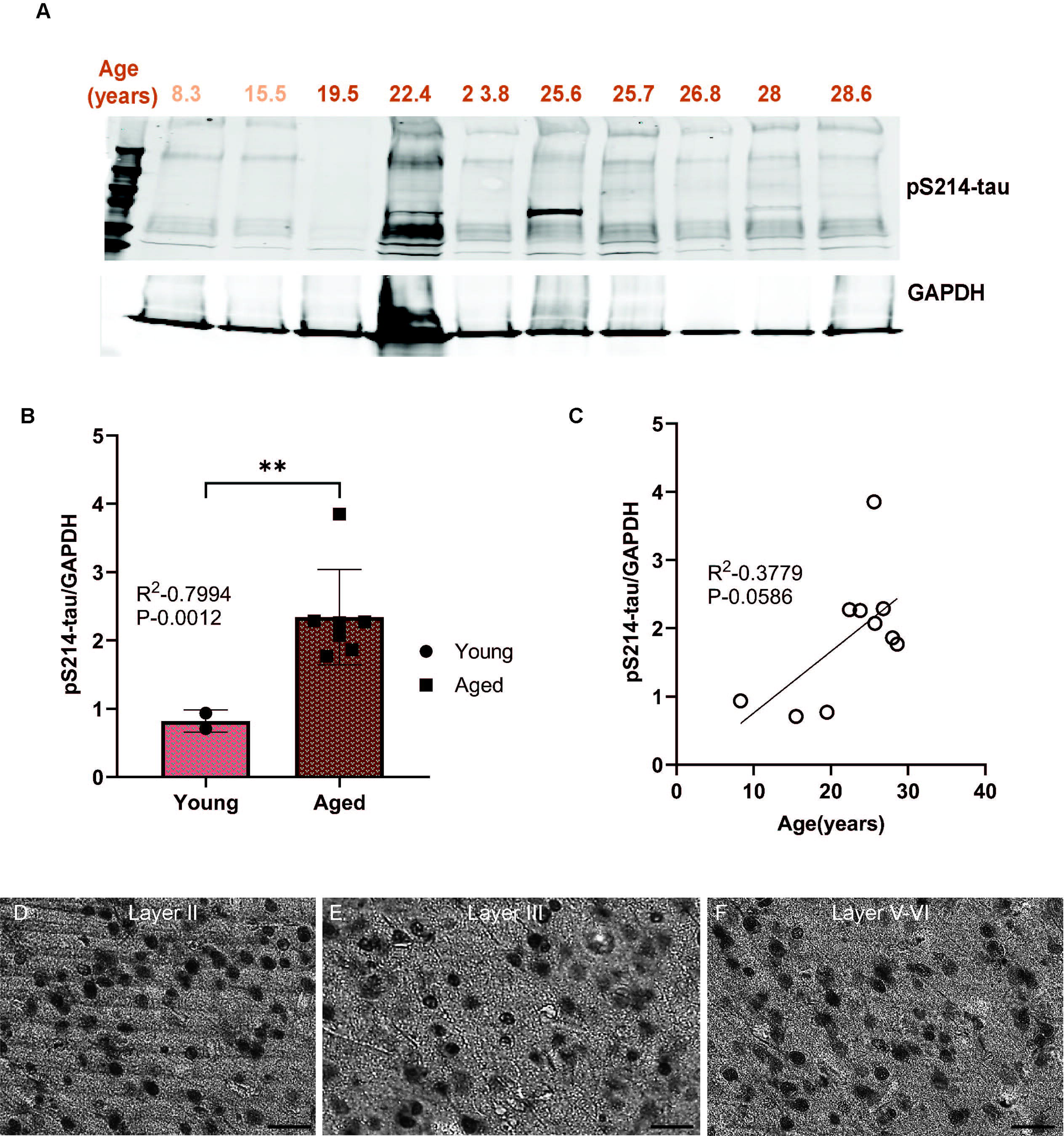
(A) Macaque ERC lysate (40 μg) was immunoblotted for pS214-tau (1:1000) and GAPDH (1:10000) across the age span (8.3-28.6 years). Animals are labeled by their age in years and color coded: young animals in light red and aged animals in dark red. (B) Expression of pS214-tau normalized by GAPDH is plotted between young (≤15.5 years) and aged (19.5-28.6 years). Young animals are denoted by circles and aged animals are denoted by squares. Means between the two groups were compared using a two-tailed unpaired *t*-test (\**P* =0.0012). (C) The correlation between levels of normalized pS214-tau and age across all animals is fit by a linear regression (R^2^ = 0.3779, \**P* = 0.0586). (D-F) Immunohistochemistry revealed robust immunolabeling for pS214-tau in aged macaque ERC, particularly in layer II, layer III and layer V-VI, within excitatory neurons along apical dendrites and in the cell body. Scale bar, 50 µm.

**Figure 2:**
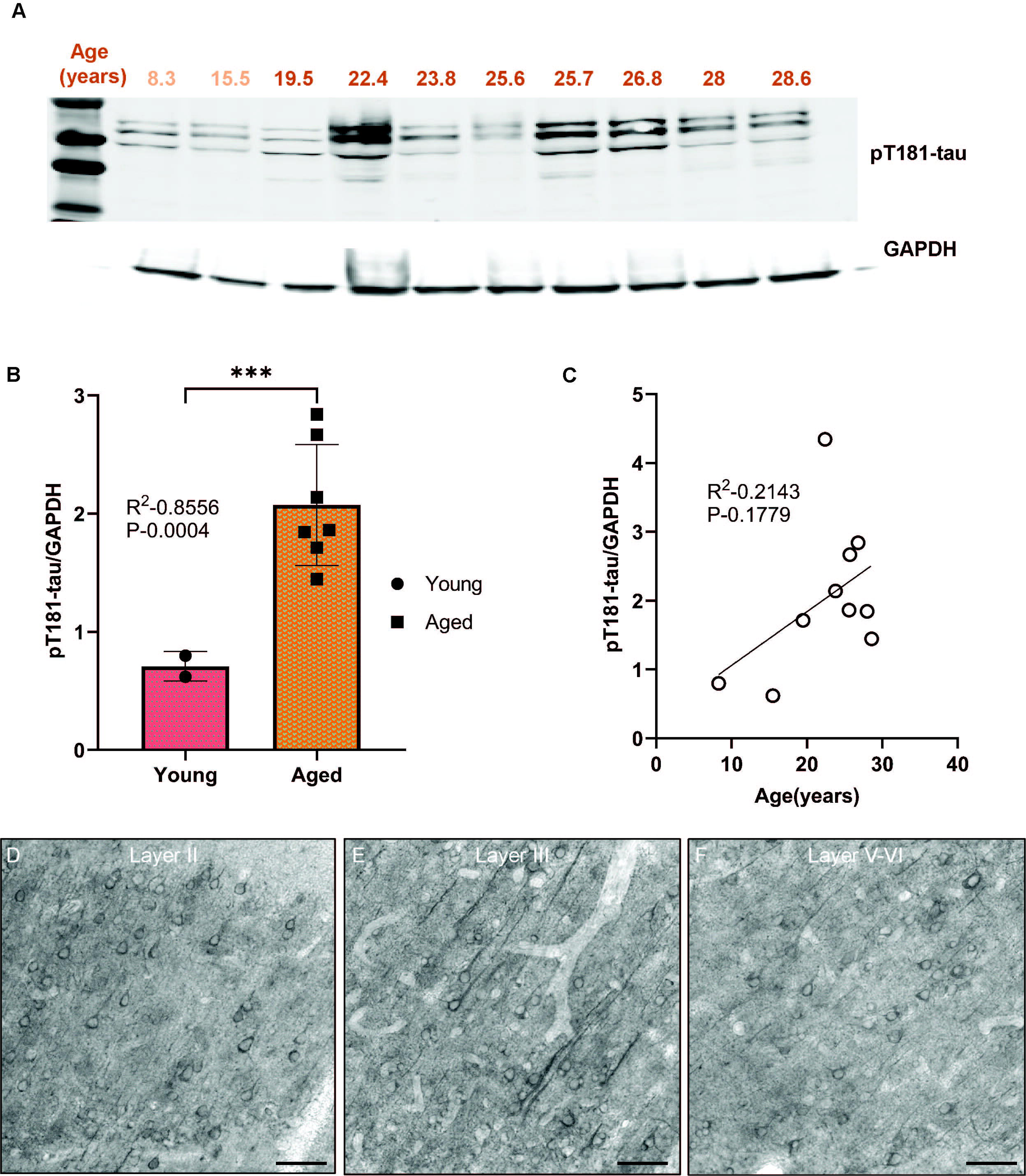
(A) Macaque ERC lysate (40 μg) was immunoblotted for pT181-tau (1:1000) and GAPDH (1:10000) across the age span (8.3-28.6 years). Animals are labeled by their age in years and color coded: young animals in light red and aged animals in dark red. (B) Expression of pT181-tau normalized by GAPDH is plotted between young (≤15.5 years) and aged (19.5-28.6 years). Young animals are denoted by circles and aged animals are denoted by squares. Means between the two groups were compared using a two-tailed unpaired *t*-test (*P =0.0004). (C) The correlation between levels of normalized pT181-tau and age across all animals is fit by a linear regression (R^2^ = 0.2143, *P = 0.1779). (D-F) Immunohistochemistry revealed immunolabeling for pT181-tau in aged macaque ERC, particularly in layer II, layer III and layer V-VI, within excitatory neurons along apical and basilar dendrites and in the cell body. Scale bar, 50 µm.

**Figure 3:**
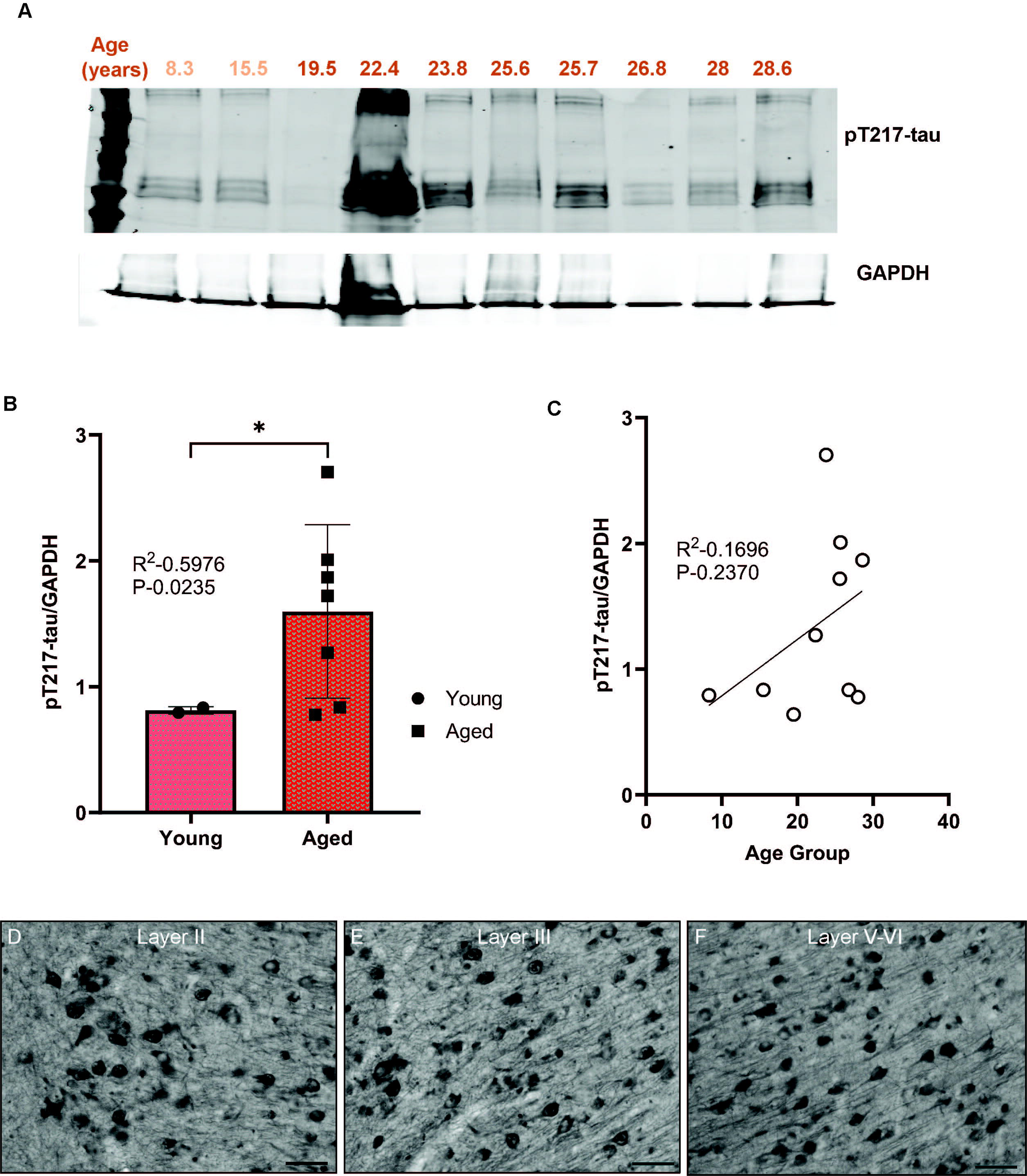
(A) Macaque ERC lysate (40 μg) was immunoblotted for pT217-tau (1:1000) and GAPDH (1:10000) across the age span (8.3-28.6 years). Animals are labeled by their age in years and color coded: young animals in light red and aged animals in dark red. (B) Expression of pT217-tau normalized by GAPDH is plotted between young (≤15.5 years) and aged (19.5-28.6 years). Young animals are denoted by circles and aged animals are denoted by squares. Means between the two groups were compared using a two-tailed unpaired *t*-test (*P =0.0235). (C) The correlation between levels of normalized pT217-tau and age across all animals is fit by a linear regression (R^2^ = 0.1696, *P = 0.2370). (D-F) Immunohistochemistry revealed robust immunolabeling for pT217-tau in aged macaque ERC, particularly in layer II, layer III and layer V-VI, within excitatory neurons along apical and basilar dendrites and in the cell body. Scale bar, 25 µm.

We used immunohistochemistry to localize pS214-tau, pT181-tau, and pT217-tau in aged macaque ERC (ages 24-31 yrs). We found robust immunolabeling for pS214-tau (**Figure 1D-F**), pT181-tau (**Figure 2D-F**), and pT217-tau (**Figure 3D-F**) phosphorylation epitopes in ERC across the cortical neuropil, including prominent labeling in stellate cells in layer II, and pyramidal cells in layer III and layers V-VI. Immunolabeling was evident in perisomatic compartments and along apical dendrites, in excitatory neurons.

### Age-related calcium dysregulation in ERC

Previous research indicated that calcium dysregulation occurs very early in the ERC (Paspalas, et al. 2018), which may help to explain why this area is the first to show cortical tau pathology. Thus, the current study examined whether there would be signs of excessive cytosolic calcium in the aged ERC, as well as evidence of reduced mGluR3 and PDE4 regulation of cAMP drive on calcium signaling.

#### pS2808-RyR2 expression in aged rhesus macaque ERC

PKA phosphorylation of RyR2 causes calcium leak from the SER into the cytosol, and has already been documented in the ERC of young adult macaques aged 7-9 years (Paspalas, et al. 2018). Consistent with previous research, there was already extensive expression of pS2808-RyR2 in macaques in this age range, with no further increase with advancing age (**Figure 4A-B**, *P =0.9307, t-0.1353, df-4.083), and no correlation between age and pS2808-RyR2 levels (**Figure 4C**, *P = 0.8505).

**Figure 4:**
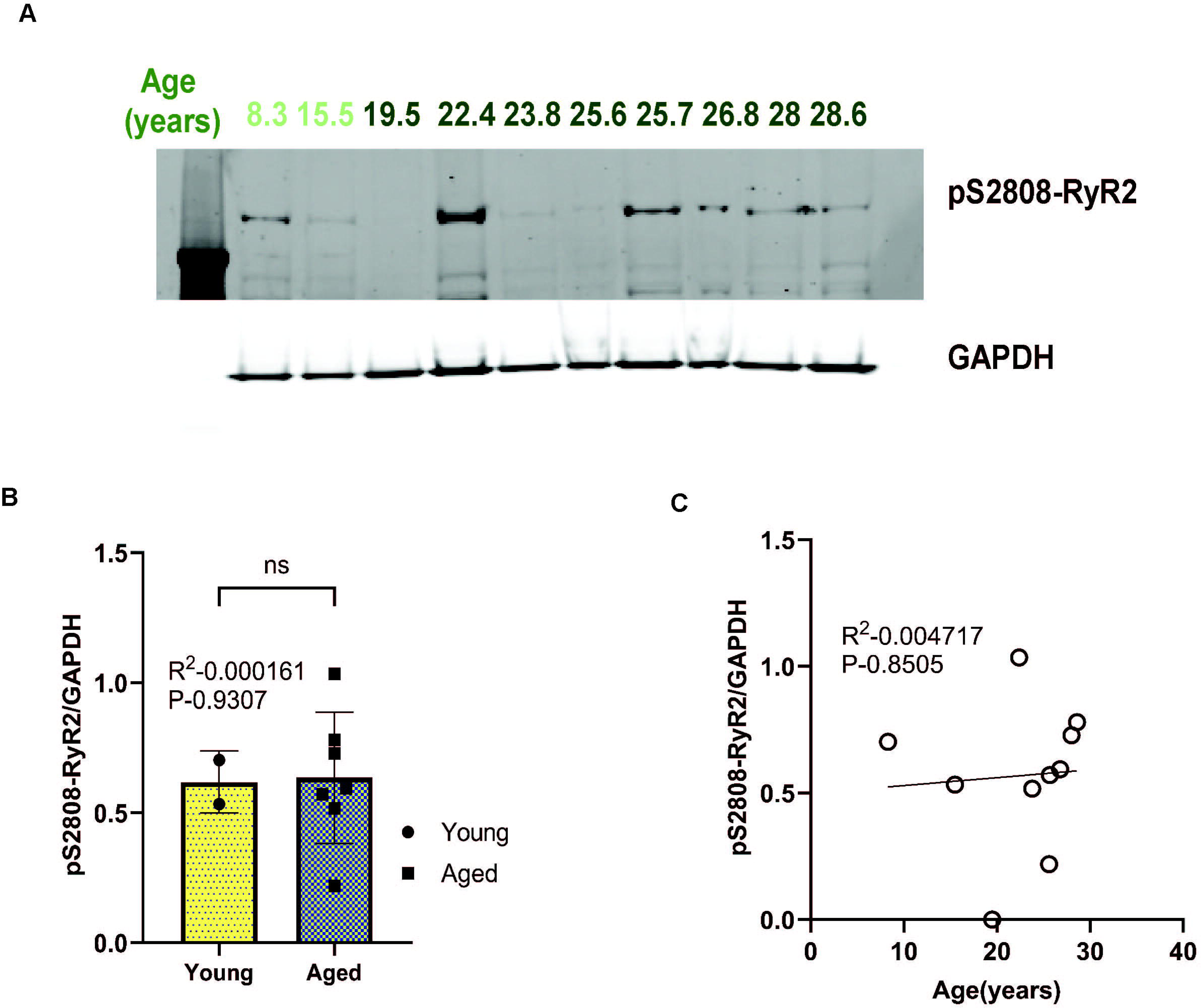
(A) Macaque ERC lysate (40 μg) was immunoblotted for pS2808-RyR2 (1:500) and GAPDH (1:10000) across the age span (8.3-28.6 years). Animals are labeled by their age in years and color coded: young animals in light green and aged animals in dark green. (B) Expression of pS2808-RyR2 normalized by GAPDH is plotted between young (≤15.5 years) and aged (19.5-28.6 years). Young animals are denoted by circles and aged animals are denoted by squares. Means between the two groups were compared using a two-tailed unpaired *t*-test (*P =0.9307). (C) The correlation between levels of normalized pS2808-RyR2 and age across all animals is fit by a linear regression (R^2^ = 0.04717, *P = 0.8505)

#### Increase in calpain 2 expression in aged rhesus macaque ERC

In contrast to calpain-1, which is activated by normal physiological levels of calcium, calpain-2 is activated by very high levels of cytosolic calcium (Baudry, et al. 2013, Higuchi et al. 2012) and can cleave and activate the kinases that hyperphosphorylate tau (Goni-Oliver et al. 2007). The expression of calpain-2 was significantly higher in aged macaque ERC than in young animals (**Figure 5A-B**, *P =0.0022, t-5.188, df-5.834), and there was a trend for a positive correlation with calpain-2 levels and age in rhesus macaque ERC (**Figure 5C**, *P =0.0947). In aged macaque ERC, using brightfield microscopy we observed calpain-2 immunolabeling in layer II stellate cell islands (**Figure 5D**), as well as in pyramidal cells in layer III (**Figure 5E**), and layer V-VI (**Figure 5F**), often expressed within apical dendrites with a twisted morphology, e.g., calpain-2 immunolabeling in layer V-VI (**Figure 5F**), common in neurofibrillary tangles (**Figure 5D-F**).

**Figure 5:**
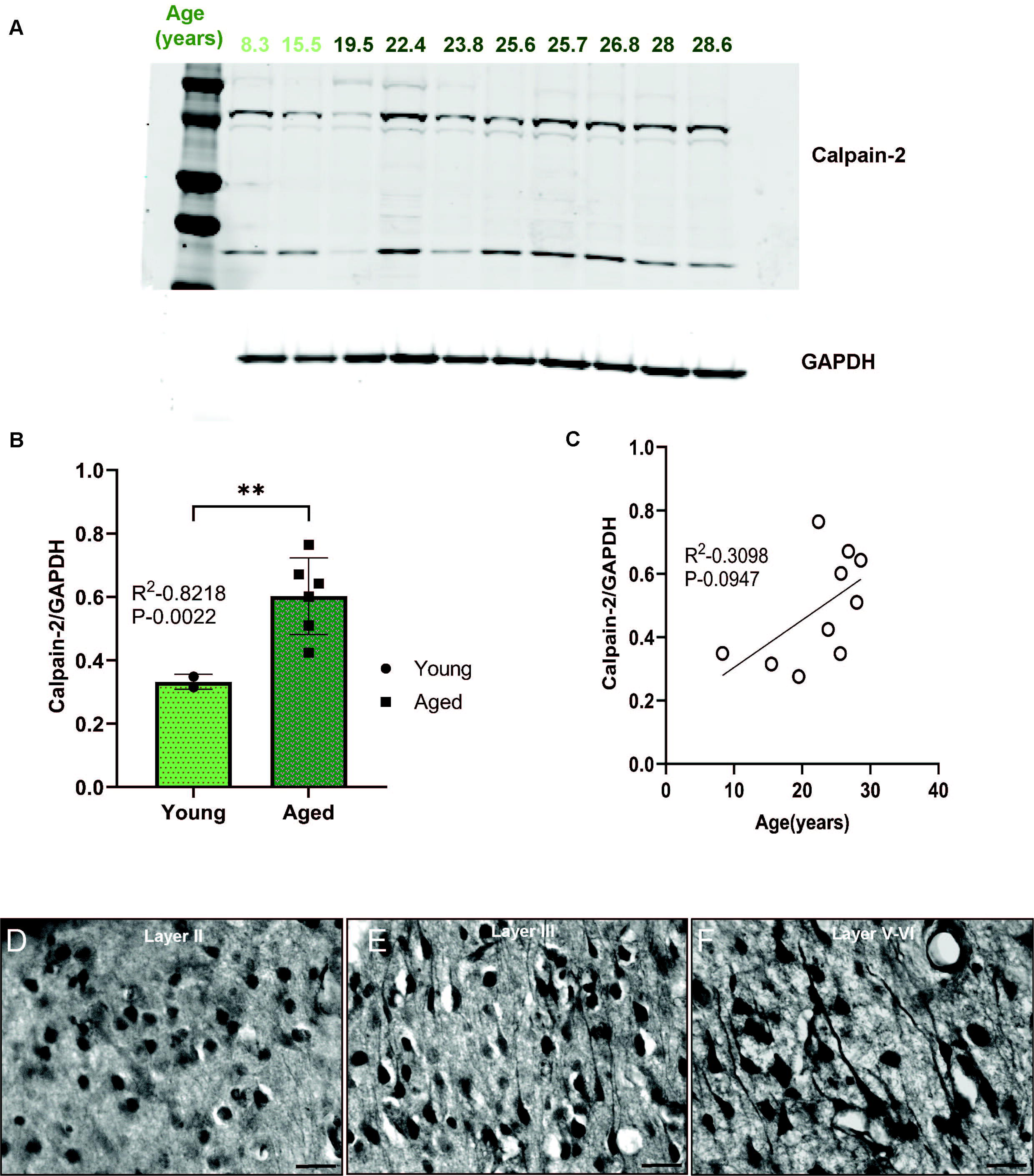
(A) Macaque ERC lysate (40 μg) was immunoblotted for calpain-2 (1:1000) and GAPDH (1:10000) across the age span (8.3-28.6 years). Animals are labeled by their age in years and color coded: young animals in light green and aged animals in dark green. (B) Expression of calpain-2 normalized by GAPDH is plotted between young (≤15.5 years) and aged (19.5-28.6 years). Young animals are denoted by circles and aged animals are denoted by squares. Means between the two groups were compared using a two-tailed unpaired *t*-test (*P =0.0022). (C) The correlation between levels of normalized calpain-2 and age across all animals is fit by a linear regression (R^2^ = 0.3098, *P = 0.0947). (D-F) Immunohistochemistry revealed robust immunolabeling for calpain-2 in aged macaque ERC, particularly in layer II, layer III and layer V-VI, within excitatory neurons along apical and basilar dendrites and in the cell body. Scale bar, 25 µm.

#### Reduced Phosphodiesterase expression in aged rhesus macaque ERC

In young adult macaques, phosphodiesterases PDE4A and PDE4D are localized on the SER, positioned to regulate cAMP-PKA drive on internal calcium release, with PDE4A generally limited to dendritic spines, and PDE4D more widely expressed with significant expression in dendrites (Carlyle et al. 2014, Datta, et al. 2020b, Datta, et al. 2021). In the current study, we found that PDE4D expression in macaque ERC decreased significantly with age (**Figure 6A-B**, *P =0.0252, t-4.967, df-2.437), while there was a modest non-significant reduction in PDE4A (**Figure 6D-E**, *P =0.5967, t-0.7210, df-1.060) expression in macaque ERC. Furthermore, there was a highly significant negative correlation between PDE4D expression and age (**Figure 6C**, *P = 0.0035). In contrast, expression of PDE4A was not correlated with age (**Figure 6F**, *P = 0.4243).

**Figure 6:**
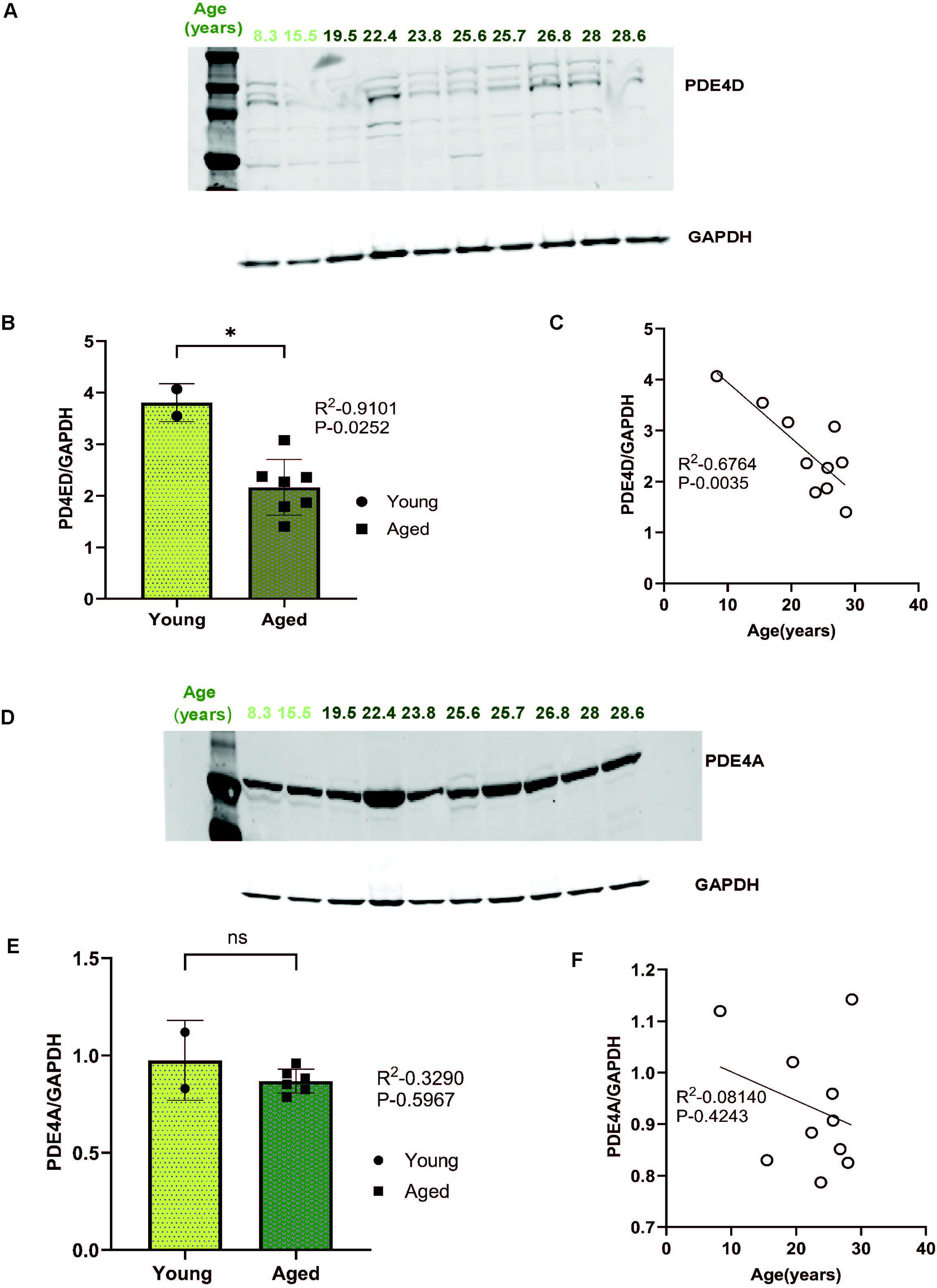
(A,D) Macaque ERC lysate (40 μg) was immunoblotted for PDE4D (1:1000), PDE4A (1:1000) and GAPDH (1:10000) across the age span (8.3-28.6 years). Animals are labeled by their age in years and color coded: young animals in in light green and aged animals in dark green. (B,E) Expression of PDE4D & PDE4A normalized by GAPDH is plotted between young (≤15.5 years) and aged (19.5-28.6 years). Young animals are denoted by circles and aged animals are denoted by squares. Means between the two groups were compared using a two-tailed unpaired *t*-test (*P = 0.0252, *P = 0.5967) respectively. (C,F) The correlation between levels of normalized PDE4D & PDE4A and age across all animals is fit by a linear regression (R^2^ = 0.6764, *P = 0.0035 & R^2^ = 0.08140, *P = 0.4243), respectively.

#### mGluR3 and GCPII expression of in aged rhesus macaque ERC

Post-synaptic mGluR3 in ERC are positioned to regulate cAMP drive on internal calcium release, a process that is reduced under inflammatory conditions by GCPII catabolism of NAAG, the endogenous ligand for mGluR3 (Datta et al. 2023). mGluR3 protein levels in aged macaque ERCs did not show significant changes in the current investigation (**Figure 7A-B**, *P =0.3629, t-1.388, df-1.243), and levels of mGluR3 were not correlated across age-span (**Figure 7C**, *P =0.5963). However, GCPII levels did increase with age (**Figure7D-E**, *P =0.0296, t-2.659, df-7) and correlated across age-span (**Figure 7F**, *P =0.2647).

**Figure 7:**
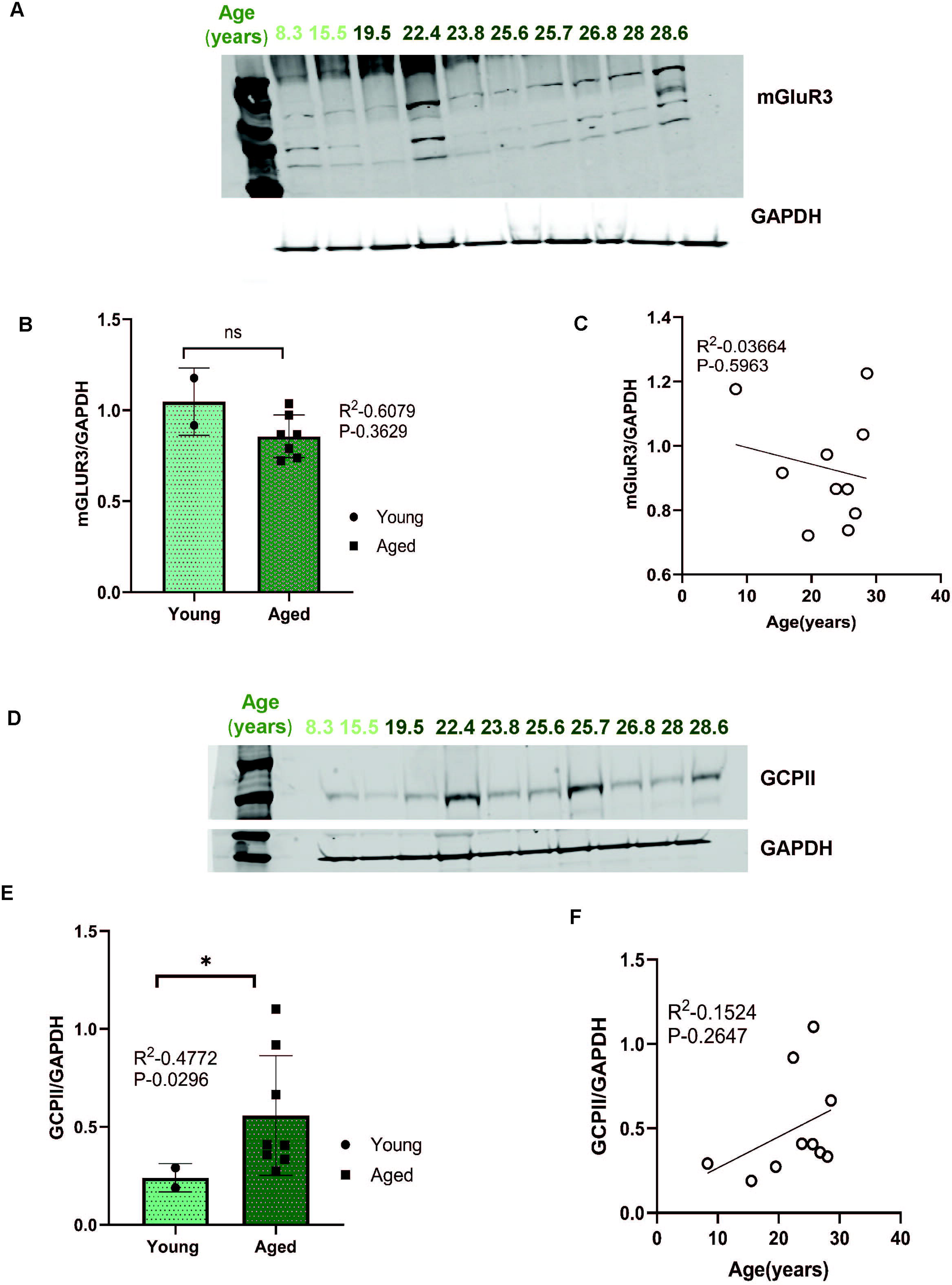
(A, D) Macaque ERC lysate (40 μg) was immunoblotted for mGluR3 (1:1000), GCPII (1:1000), and GAPDH (1:10000) across the age span (8.3-28.6 years). Animals are labeled by their age in years and color coded, young animals in in light green and aged animals in dark green. (B, E) Expression of mGluR3 and GCPII normalized by GAPDH is plotted between young (≤15.5 years) and aged (19.5-28.6 years) age respectively. Young animals are denoted by circle and aged animals are denoted by square. Means between the two groups were compared using a two-tailed unpaired *t*-test (*P =0.3629 & *P = 0.9307). (C) The correlation between levels of normalized mGluR3, GCPII and age across all animals is fit by a linear regression (R^2^ = 0.03664, *P =0.5963, & R^2^ = 0.1524, *P = 0.2647) respectively.

### Correlations between evidence of dysregulated calcium signaling and tau hyperphosphorylation

We examined potential correlations between measures of cAMP-calcium dysregulation and tau hyperphosphorylation. We found that PDE4D expression had an inverse correlation with pTau levels, where reduced levels of PDE4D significantly correlated with increased levels of pS214-tau (**Figure 8A**, *P=0.0383), consistent with dysregulated cAMP signaling increasing tau phosphorylation by PKA at S214 (Carlyle, et al. 2014, Datta, et al. 2021). Reduced levels of PDE4D also correlated with increased levels of pT217-tau (**Figure 8B**, *P =0.0111), but showed only a trend level inverse correlation with pT181-tau (p=0.3).

**Figure 8:**
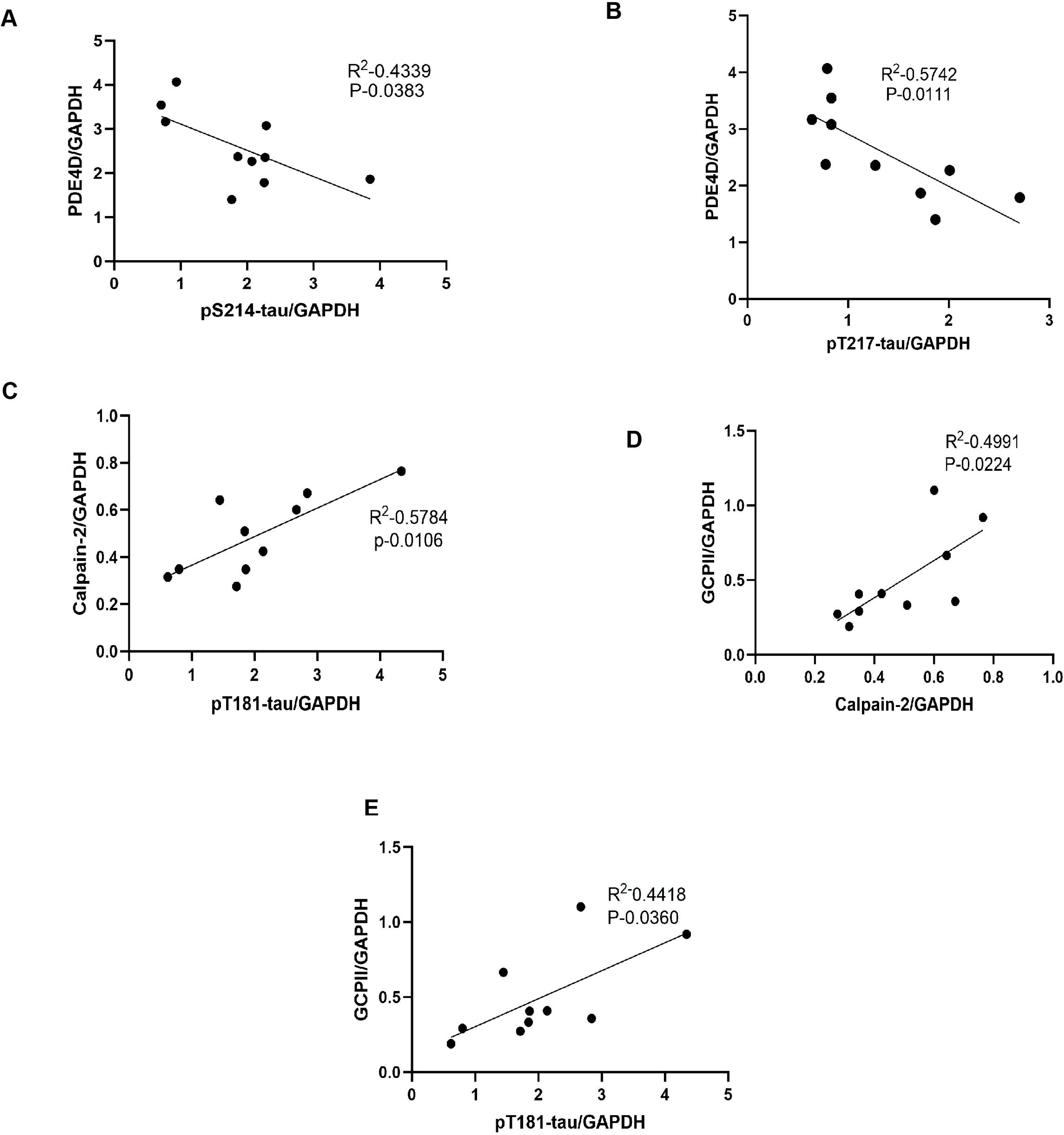
(A) Correlation between levels of pS214-tau by GAPDH (x axis) and PDE4D by GAPDH (y axis) is fit by a linear regression (R^2^ = 0.4339, *P-0.0383). (B) Correlation between levels of pT217-tau by GAPDH (x axis) and PDE4D by GAPDH (y axis) is fit by a linear regression (R^2^ = 0.5742, *P-0.0111). (C) Correlation between levels of pT181-tau by GAPDH (x axis) and calpain-2 by GAPDH (y axis) is fit by a linear regression (R^2^ = 0.5784, *P = 0.0106). (D) Correlation between levels of calpain2 by GAPDH (x axis) and GCPII by GAPDH (y axis) is fit by a linear regression (R^2^ = 0.4991, *P-0.0224). (E) Correlation between levels of pT181-tau by GAPDH (x axis) and GCPII by GAPDH (y axis) is fit by a linear regression (R^2^ = 0.4418, *P-0.0360).

Conversely, we found a strong positive correlation between calpain-2 levels and pT181-tau expression (**Figure 8C**, *P = 0.0106), while the positive correlations between calpain-2 levels and pS214-tau and pT217-tau levels did not reach significance (p=0.38 and 0.6, respectively). There was also a positive correlation between GCPII levels and calpain-2 levels, consistent with reduced mGluR3 regulation of cAMP drive on calcium signaling (**Figure 8D**, *P = 0.0224). GCPII expression generally correlated with pTau levels, with pT181-Tau levels reaching statistical significance (**Figure 8E**, *P = 0.0360). Correlations with pT217-tau and pS214-tau showed a trend but did not reach significance due to a single animal with high levels of tau, but intermediate levels of GCPII, consistent with multiple factors contributing to tau hyperphosphorylation.

## Discussion

The current study found increases in phosphorylated tau (pT181-tau, pS214-tau, pT217-tau) and evidence of increased inflammation (GCPII) and calcium dysregulation (calpain-2) in the aged macaque ERC. There was a positive correlation between GCPII and calpain-2 levels, consistent with GCPII dysregulating calcium signaling, and general positive correlations between levels of GCPII, calpain and pTau species. Conversely, there were negative correlations between levels of PDE4D and pTau species. Overall, these data indicate an environment of dysregulated cAMP-calcium signaling in the aging ERC that is associated with the rise of early stage, soluble phosphorylated tau. At this early stage, soluble pTau is likely the form that traffics between neurons to seed tau pathology in a network of excitatory neurons (Datta et al. 2024), therefore, characterization at this early stage is particularly important for developing informed strategies for disease prevention. The increase in GCPII inflammation, and its correlations with calpain-2 and pTau levels are of special interest given the likely roles of inflammatory mechanisms in the common, sporadic form of AD, and the previous finding that levels of pT217-tau correlate with GCPII activity in the dlPFC (Bathla et al. 2023). Altogether, these data suggest that targeting inflammation, and specifically calcium dysregulation, may be beneficial in reducing early tau pathology.

A weakness of the current study is the relatively small number of subjects, due in large part to the current scarcity of macaques in general given their extensive use in creating SARS-CoV-2 vaccines, and the further rarity of macaques that reach very old age. In particular, larger numbers of subjects may have allowed trends in correlation with age to be significant. It is noteworthy that this type of work cannot be done in rodent models that depend on autosomal dominant mutations rather than inflammation to cause pathology, nor in humans, where PMI longer than 15 min precludes opportunity to capture soluble pTau (Matsuo, et al. 1994, Wang, et al. 2015). Thus, the current data is very valuable for revealing early molecular events in primate ERC related to the rise in tau pathology.

### Increased pTau with age in ERC

Assays of the aging macaque ERC documented elevated phosphorylated tau, consistent with this region being the earliest site of cortical tau pathology. Previous data had shown evidence of increasing hyperphosphorylation of tau with age in the ERC with both increasing molecular weights and increasing insolubility (Paspalas, et al. 2018). The current study builds on this by documenting the rise in pT181, pS214 and pT217. These are all early tau phosphorylation sites (Wesseling et al. 2020), with pT217-tau and pT181-tau emerging as important fluid biomarkers of AD (Barthelemy et al. 2023, Horie et al. 2023, Janelidze et al. 2021).

Plasma pT217-tau in particular is revolutionizing the field, as it heralds future disease, and consistently discriminates AD from other neurodegenerative diseases, appears in the earliest presymptomatic stages of AD, and correlates strongly with premortem neuropathological tau burden (Barthelemy, et al. 2020, Hansson, et al. 2023, Janelidze, et al. 2020, Mattsson-Carlgren, et al. 2023, Olsson, et al. 2016, Salvado, et al. 2023). Recent data show that the rise in plasma pT217-tau correlates especially well with the appearance of Aβ PET signals in brain (Ashton et al. 2024, Barthelemy, et al. 2023, Barthelemy et al. 2024, Janelidze et al. 2024, Janelidze, et al. 2021), but little has been known about its rise in brain. Our recent immunoEM studies have documented aggregations of pT217-tau in the dendrites and dendritic spines of layer II ERC neurons from “early” aged (18-19 years) macaques (Datta, et al. 2024), consistent with the current biochemical findings. Nanoscale imaging also demonstrated pT217-tau trafficking between synapses, where it can be captured in extracellular fluid (Datta, et al. 2024), helping to explain how this pTau species reaches CSF and plasma as a fluid biomarker.

### Support for dysregulated cAMP-calcium inflammatory signaling with age

Decades of research indicate that calcium dysregulation is an early driver of pathology in both sporadic and autosomal dominant disease (Alzheimer’s Association Calcium Hypothesis 2017, Gant et al. 2018, Khachaturian 1994), with elevated calcium in the cytosol, rather than stored in the SER, activating calpain-2 to disinhibit GSK3β and cdk5 to hyperphosphorylate tau (Arnsten, Datta and Preuss 2021, Arnsten, Datta and Wang 2021). cAMP-PKA calcium signaling plays an important role in driving calcium release out of the SER, which is regulated by PDE4D anchored to the SER to reduce cAMP-PKA signaling (Bathla, et al. 2023). The current study showed a loss of PDE4D in the aged ERC, suggesting disrupted regulation of cAMP-PKA signaling, leading to excessive PKA phosphorylation of RyR2 (pRyR2) causing calcium to “leak” into the cytosol from the SER. The current study replicated earlier data showing that pRyR2 expression is already evident in the ERC in middle age, and additionally documented increased expression of calpain-2, which plays an important role in driving both tau hyperphosphorylation and autophagic degeneration, and is associated with neurofibrillary tangles in AD brains (Adamec et al. 2002, Grynspan et al. 1997). Decreased PDE4D levels and increased calpain-2 levels correlated with increased pTau levels in the aged macaque ERC, consistent with dysregulated cAMP-calcium signaling contributing to tau hyperphosphorylation.

### mGluR3 intact as potential therapeutic target: Focus on the role of GCPII inhibitors

While mGluR3 have traditionally been considered presynaptic receptors based on their localization in rodent (Woo et al. 2022), new data show that mGluR3 have a very different and important role in primate higher cortical circuits, where they are post-synaptic and regulate cAMP-calcium signaling (Arnsten and Wang 2020). This has been seen in both dlPFC (Jin et al. 2018a) and more recently in the macaque ERC (Datta, et al. 2023). mGluR3 are stimulated not only by glutamate, but by NAAG which is selective for mGluR3 (Neale and Olszewski 2019). However, NAAG-mGluR3 signaling is a target of inflammation when GCPII catabolizes NAAG (Arteaga Cabeza et al. 2021, Zhang et al. 2016). The current study found an age-related increase in GCPII in the macaque ERC which correlated with increased pTau expression. These data are consistent with previous work showing that chronic GCPII inhibition in aged monkeys reduces pT217-tau levels in the ERC and dlPFC, as well as in blood (Bathla, et al. 2023). This study also found a strong correlation between levels of GCPII activity and pT217-tau expression in dlPFC (Bathla, et al. 2023), emphasizing the relevance of this pathway to early-stage AD pathology. As the current study found that mGluR3 expression remains relatively intact, this beneficial substrate appears to remain in dlPFC, further encouraging the development of GCPII inhibitors for human trials.

## Acknowledgements

We thank Caroline Zeiss, Dan Holden, Lisa Ciavarella, Tracy Sadlon, Sam Johnson, and Michelle Wilson for their technical expertise and assistance with animals.

